# The Latency of a Domain-General Visual Surprise Signal is Attribute Dependent

**DOI:** 10.1101/2025.08.18.670829

**Authors:** Benjamin G. Lowe, Naohide Yamamoto, Jonathan Robinson, Patrick Johnston

## Abstract

Predictions concerning upcoming visual input play a key role in resolving percepts. Sometimes input is surprising, under which circumstances the brain must calibrate erroneous predictions so that perception is veridical. Despite the extensive literature investigating the nature of prediction error signalling, it is still unclear how this process interacts with the functionally segregated nature of the visual cortex, particularly within the temporal domain. Here, we recorded electroencephalography (EEG) from humans whilst they viewed static image trajectories containing a bound object that sequentially changed along different visual attribute dimensions (shape and colour). Crucially, the context of this change was designed to appear random (and unsurprising) or violate the established trajectory (and cause a surprise). Event-related potential analysis found no effects of surprise after controlling for cortical adaptation. However, multivariate pattern analyses found whole-brain neural representations of visual surprise that overlapped between attributes, albeit at distinct, attribute-specific latencies. These findings suggest that visual surprise results in whole-brain, generalised (i.e., attribute-agnostic) prediction error responses that conform to an attribute-dependent temporal hierarchy.

## Introduction

Predictive coding postulates that the brain uses prior knowledge to predict visual events, with said predictions being integrated with upcoming input on a moment-by-moment basis (Friston, 2005, 2010). Of course, sometimes visual inputs violate our expectations (e.g., when your friend surprises you with a new and daring hairstyle), which, according to theory, causes a cascade of ‘prediction error’ signalling that recalibrates the relevant erroneous predictions generated by the brain. To date, the mechanisms through which our brains signal prediction error are still hotly debated (e.g., Feuerriegel et al., 2021; Press et al., 2020; Walsh et al., 2020; Westerberg et al., 2025), with most work skirting around how this process might depend on the specific stimulus attributes (or ‘features’) that violate an expectation.

Entertaining this nuance gives rise to a myriad of interesting questions, given the functionally segregated nature of the visual system. For instance, it is well established that the different attributes comprising a visual object are processed within distinct regions of cortex (Barrett et al., 2001; Goodale et al., 1991; Haxby et al., 1991; Mishkin et al., 1983; Schneider, 1969; Seymour et al., 2010; Zeki et al., 1991), at different relative latencies (Arnold, 2005; Clifford et al., 2003, 2004; Holcombe, 2009; Moutoussis & Zeki, 1997b, 1997a). This strongly suggests that the nature of a prediction error signalling ought to depend on the specific attribute violating an expectation. To date, only a small handful of studies have begun to address this empirically (Jiang et al., 2016; Robinson, Woods, et al., 2020; Stefanics et al., 2018, 2019), with said work being principally concerned with identifying the spatial origins of these signals. Thus, it is unclear how attribute-specific prediction error might differ within the temporal domain.

Recently, we demonstrated that surprising inputs from different attributes evoke overlapping neural responses, albeit at distinct latencies (Lowe et al., 2023). Specifically, electroencephalographic (EEG) responses evoked by violating a size expectation were faster, but otherwise shared, with those evoked by violating an orientation expectation within the same stimulus. This suggests the existence of a temporal hierarchy in prediction error signalling, and aligns well with work showing that surprise evoked from one attribute ‘spreads’ across all other areas involved in encoding the corresponding stimulus (Jiang et al., 2016). Collectively, these works assert that an entire bound object is rendered unexpected if surprise is signalled for just one of its attributes, with the latency of this process depending on which attribute was surprising.

While we find these results compelling, the sequential manner by which we manipulated both size (increasing vs. decreasing trajectories) and orientation (clockwise vs. anti-clockwise trajectories) may have been perceived as apparent motion. Thus, the inferred overlap in neural representation between violation types could have reflected a higher-order prediction error signal concerning stimulus motion, rather than a generalised prediction error response evoked by independent attributes. To rule out this alternative explanation, we conducted a similar experiment, this time independently manipulating expectations concerning the shape (or ‘form’) and colour of a bound stimulus. Not only are these attributes impossible to conflate with another high-order attribute, but they are also independently encoded within the visual system (e.g., Barret et al., 2001; Clifford et al., 2004; Seymour et al., 2010).

In alignment with our previous work, we hypothesised that visual surprise signalling, evoked from shape and colour, would share spatially overlapping neural representations that are temporally distinct. This hypothesis was supported by our results, suggesting that 1) our previous findings were not due to a common higher-order attribute violation (i.e., motion extrapolation), and 2) support the notion that there is a temporal hierarchy for generalised visual surprise signalling within the visual system.

## Materials & Methods

### Participants

Thirty-five participants were recruited for the experiment, which was approved by Queensland University of Technology’s Human Research Ethics Committee (approval no. 1900000739). Inclusion criteria required that the participants had (corrected to) normal vision and no history of neurological disorder, all indicated through self-report. Signed consent for participation was collected before beginning the experiment, and participants were compensated with partial course credit or $10 (AUD). We excluded three participants who demonstrated poor vigilance during the experimental session (see Monitoring Participant Vigilance). Thus, our final sample consisted of 32 participants (22 women and 10 men), aged between 17 and 45 years (*M* = 23.71 years, *SD* = 7.96 years).

### Visual Stimulation

Participants viewed a novel contextual trajectory paradigm (Johnston et al., 2017) on a HP Liquid Crystal display (1920 x 1080 px resolution, 60 Hz refresh rate) within a dimly-lit room, from a distance of ∼58 cm. Using version 1.90.2 of PsychoPy (Peirce et al., 2019), we varied three attributes (shape, colour, and orientation) of a centrally-presented visual stimulus object (spanning ∼4.5° of visual field) to create trial-wise change sequences with varying levels of attribute-specific contextualised predictability (Figure 1). The sequences began with a 500 ms fixation cross followed by four static image ‘steps’ (500 ms per image; 0 ms inter-stimulus interval).

**Figure 1.**
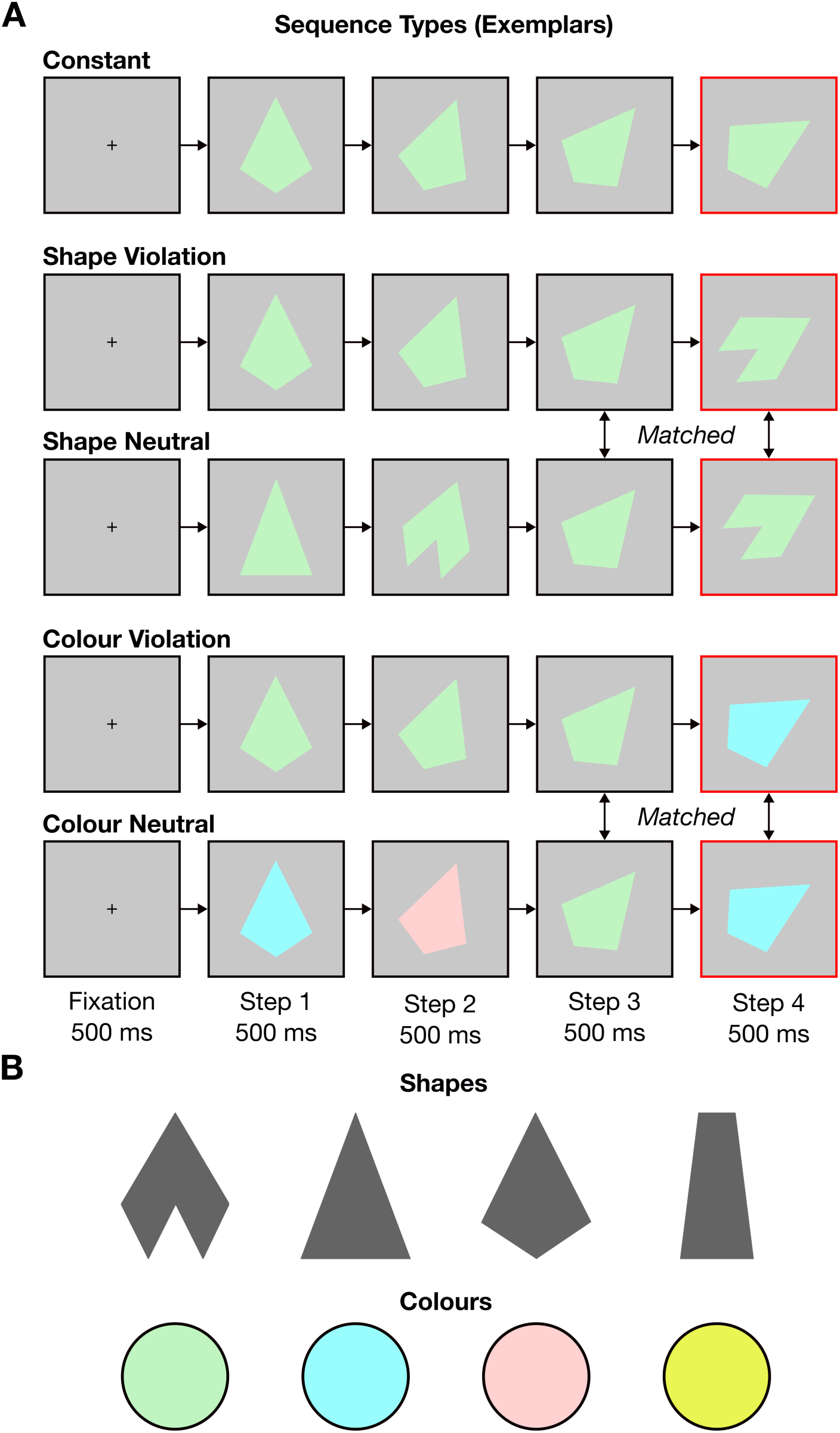
Visual stimulation paradigm. **A)** Exemplars of each sequence type (0 ms inter-stimulus interval between steps). Note that the final two steps are identical across the violation and neutral conditions within a given level of attribute. Any differences in evoked responses from Step 4 (red outline) must have therefore reflected the preceding change context. **B)** The shapes and colours used for generating sequences. Participants were shown all possible shape-colour conjunctions, during both the penultimate and final steps, an equal number of times within and between conditions. Orientation violation sequences are shown within Supplementary Materials (Figure S3A) because the corresponding data have been omitted from the main text.

The first three sequence steps were arranged to induce simultaneous expectations (or lack thereof) concerning three attributes of a fourth step that was yet to be displayed (colour, shape, and orientation). During 288 trials, the final step violated the established trajectory for one of the three attributes (balanced across shape, colour, and orientation), serving as surprising attribute-specific visual inputs. So that participants did not learn to expect trajectory violations, we included an equal number of constant sequence trials, where the final step conformed to the established trajectory. Thus, there was a 50% chance of a violation trial if the first three sequence images established a coherent trajectory. Finally, there were 192 neutral sequence trials, where one of the attributes varied unpredictably in each step (balanced across shape and colour). These were intended to evoke feedforward attribute-specific neural activity that was largely devoid of experimentally induced expectations. Participants took self-paced breaks every ∼170 trials to reduce fatigue (four breaks total).

A critical aspect of our design was that the set of transition exemplars between sequence steps 3 and 4 were identical across violation and neutral trials within each level of attribute (Figure 1A). As such, any differences in evoked responses must have reflected the contextually induced levels of surprise, and not the physical properties of stimuli themselves. Moreover, because the attribute in these trials always *changed* relative to that shown immediately beforehand, differences in evoked responses were highly unlikely to be confounded by a release from cortical adaptation (e.g., repetition suppression). This is otherwise a common critique of seemingly similar oddball-type designs, where violations (‘deviants’) are compared to repeatedly presented stimuli (Amado & Kovács, 2016; Feuerriegel et al., 2021; Solomon et al., 2021; Westerberg et al., 2025).

Note that findings concerning orientation have been omitted from the main text because this attribute lacked a neutral change control condition. We intended to compare violations here to the constant condition; however, unlike with the other attributes, this approach would have failed to isolate the effects of surprise from those of a confirmed expectation. To elaborate, responses to surprising and expected input are increasingly becoming viewed as distinct effects within the contemporary literature, rather than two sides of the same coin (e.g., Feuerriegel et al., 2021; Grotheer & Kovács, 2016; Press et al., 2020). Thus, isolating the effects of visual surprise requires a control condition devoid of experimentally induced expectations (e.g., the neutral change sequences for shape and colour in Figure 1A). For completeness, results for orientation are reported in our Supplementary Materials (Figure S3-Figure S4).

### Monitoring Participant Vigilance

We used rarely occurring catch trials to probe whether participants remained vigilant throughout the experimental session. These were made by randomly cloning a subset of 77 trials within the trial list (9.11% of the total trial count, including catch trials) wherein a grey dot probe onset on top of the stimulus during one of the first three steps within a trial. Participants needed to respond to each probe before the end of the trial sequence using the spacebar on a keyboard. Those who failed to detect at least 90% of these probes were rejected (*n* = 2). Catch trials were excluded from analyses.

### EEG Acquisition and Preprocessing

We used a 64-channel BioSemi ActiveTwo system to record EEG at a sampling rate of 1024 Hz with ActiView software (version 7.06, Biosemi). Channels were laid out using the 10-10 international placement system, with the common mode sense (CMS) as an online reference and driven right leg (DRL) serving to reduce common mode interference and improve signal quality.

We preprocessed the data offline using version 1.6 of MNE-Python (Gramfort et al., 2013) within a Python 3.9.16 environment. Here, we created two sets of preprocessed data per participant that were used for univariate event-related potential (ERP) and multivariate decoding analyses, respectively. The ERP data were preprocessed between 1 and 30 Hz, while the decoding data were preprocessed between 0.1 and 100 Hz—all subsequent preprocessing steps were identical between data sets. The data were notch filtered at 50 Hz, and excessively noisy channels were removed and interpolated (*M* = 0.25 interpolations across participants; *SD* = 0.66). We then re-referenced the data to the average activity across the whole scalp and created epochs between -2 and 1 s relative to the final step’s onset within each sequence (encapsulating the entire trial sequence). It is important to note that our event triggers onset two frames before each stimulus onset, but we accounted for this before segmenting the data into epochs (Figure S1). Finally, we downsampled the data to 250 Hz, reducing the computational cost of subsequent analyses, and then baseline-corrected each epoch using the mean activity between -100 and 0 ms relative to the final step onset.

### ERP Component Analysis

Like in previous work (Baker et al., 2021, 2022; Johnston et al., 2017; Robinson, Breakspear, et al., 2020), we first compared whether ERP responses evoked from the final image step differed between conditions. Specifically, we extracted each participant’s peak amplitude value within the P1 (75 to 125 ms post stimulus onset) and N170 (150 to 250 ms post stimulus onset), averaged over occipito-parietal electrodes (O1, O2, PO7, PO8, P7, P8, P9, and P10 as per Robinson, Breakspear, et al. 2020). We then separately subtracted component peaks evoked by the constant sequences from the attribute-specific change context sequences, and compared the resulting differences to zero using one-tailed Bayesian *t*-tests (half-Cauchy prior; *r* = 0.707) in version 0.18.3 of JASP (JASP Team, 2024).

In short, these quantified the ratio of evidence favouring the alternative hypothesis compared to the null hypothesis in the form of a Bayes factor (BF_10_; Rouder et al., 2009). BF_10_ values greater than 1 provide evidence for an effect, and those smaller than 1 provide evidence for no effect. BF_10_ values between 1/3 and 3 were deemed to offer inclusive evidence for either hypothesis. Note that the tests were one-tailed because constant sequences were hypothesised to evoke smaller responses (in absolute terms) than those evoked by the change sequences, due to the former being subject to repetition suppression at the attribute level (Amado & Kovács, 2016; Feuerriegel, 2024; Feuerriegel et al., 2021; Solomon et al., 2021; Westerberg et al., 2025).

Next, we compared the peak responses between change sequence types. This was done separately for the P1 and N170, using 2 x 2 Bayesian repeated measures analyses of variance (ANOVAs) with attribute (shape vs. colour) and change context (neutral vs. violation) as factors. Briefly, this analysis quantifies how much more likely the data are under models including each factor, or the interaction term, compared to models that omit them, in the form of a Bayes factor (BF_incl_; Rouder et al., 2012). As with BF_10_ values, BF_incl_ values above one provide evidence for an effect and values below one provide evidence for no effect. BF_incl_ values between 1/3 and 3 were interpreted to be inconclusive for determining whether the factors contributed to the models. We hypothesised a main effect whereby violation sequences would evoke greater neural activity (in absolute terms) compared to the neutral sequences by virtue of increased prediction error signalling (Friston, 2005, 2010).

### Decoding Analyses

Our main research question concerned whether 1) aspects of a visual surprise signal were shared between independent attributes within a bound stimulus object (here shape and colour); and 2) whether the latency of this overlap differed between attributes (Lowe et al., 2023). To this end, we employed linear discriminant analysis (LDA) classifiers to firstly decode change context, separately for shape and colour, using multivariate patterns of activity evoked from neutral vs. violation trials following the fourth image step (Figure 1A). Crucially, this approach was cross-validated in a leave-one-exemplar-out manner such that all physical stimulus characteristics were balanced between conditions within both the training and test folds (96 folds per comparison). Thus, any decoded signal must have reflected the context that a stimulus was shown within and not the stimulus itself. We further conducted a qualitative searchlight analysis at each time point (searchlight size of five electrodes) to coarsely localise where on the scalp any decoded signal might have originated from (Kriegeskorte et al., 2006).

After separately decoding surprise for shape and colour, we cross-decoded this signal between attributes. Here, we trained LDA classifiers to decode change context from all shape neutral and shape violation trials, and then used the resulting decision rules to classify change context from colour neutral and colour violation trials (and vice versa). Because the physical stimulus characteristics were identical across conditions within an attribute comparison, but different from the stimuli used within the other attribute comparison (see Figure 1A), any above-chance decoding must have reflected surprise signalling that generalises across shape and colour attributes.

Because we were concerned with the relative latency in an otherwise shared process between attributes, this approach was temporally generalised (King & Dehaene, 2014). By this, we mean that the classifiers trained on each time point of data were generalised to all time points within the epoch, resulting in temporal generalisation matrices (TGMs). The diagonal of these TGMs corresponded to when training and test times were equivalent. Therefore, latency shifts were apparent when a signal was cross-decoded exclusively either above (i.e., training time *later* than test time) or below (i.e., training time *earlier* than test time) the diagonal. To aid our interpretation, participant-level TGMs were smoothed using a two-dimensional Gaussian filter (σ = 20 ms; see Figure S2 for unsmoothed data).

### Statistical Inference

Time point-level inference concerning decoded data was conducted using Bayesian *t*-tests within version 0.9.12-4.7 of the BayesFactor R package (Morey & Rouder, 2023). As recommended by Teichmann et al. (2022), one-tailed tests employed a half-Cauchy prior with default width (*r* = 0.707) and null interval between δ = 0 and δ = 0.5. Similarly, two-tailed tests employed a full Cauchy prior with default width and null interval between δ = -0.5 and δ = 0.5.

We additionally ran null hypothesis tests using cluster-level inference on our TGMs, which is more conventional within the neuroimaging literature (Friston et al., 1996; Maris & Oostenveld, 2007). Specifically, we generated non-parametric null distributions of maximum cluster sizes using two-tailed *t*-tests on shuffled data (5000 permutations), with a cluster-defining threshold of *p* < 0.01. Afterwards, we identified cluster sizes within our unshuffled data that were larger than the 95th percentile of the said null distribution. These were interpreted as significant at a family-wise error rate of 0.05.

## Results

### Univariate ERP Comparisons Between Trajectory Types

We first tested whether presenting identical stimuli within different contextual trajectories affected the amplitude of evoked responses. As expected, the fourth step of each condition evoked canonical P1 and N170 ERP components over occipito-parietal channels (Figure 2). Moreover, the amplitude of these components were larger during the change trajectory conditions compared to the constant trajectory condition. This was clearly the case for violated colour and violated shape during the P1 (BF_10_ = 21.54 and 55.52, respectively), and neutral shape, violated shape, and violated colour during the N170 (BF_10_ = 15.52, 4.44, and 3.16, respectively). On the other hand, Bayes factors provided inconclusive evidence for the presence or absence of an effect in the neutral shape vs. constant P1 comparison and the neutral colour vs. constant P1 and N170 comparisons (Figure 2B-C).

**Figure 2.**
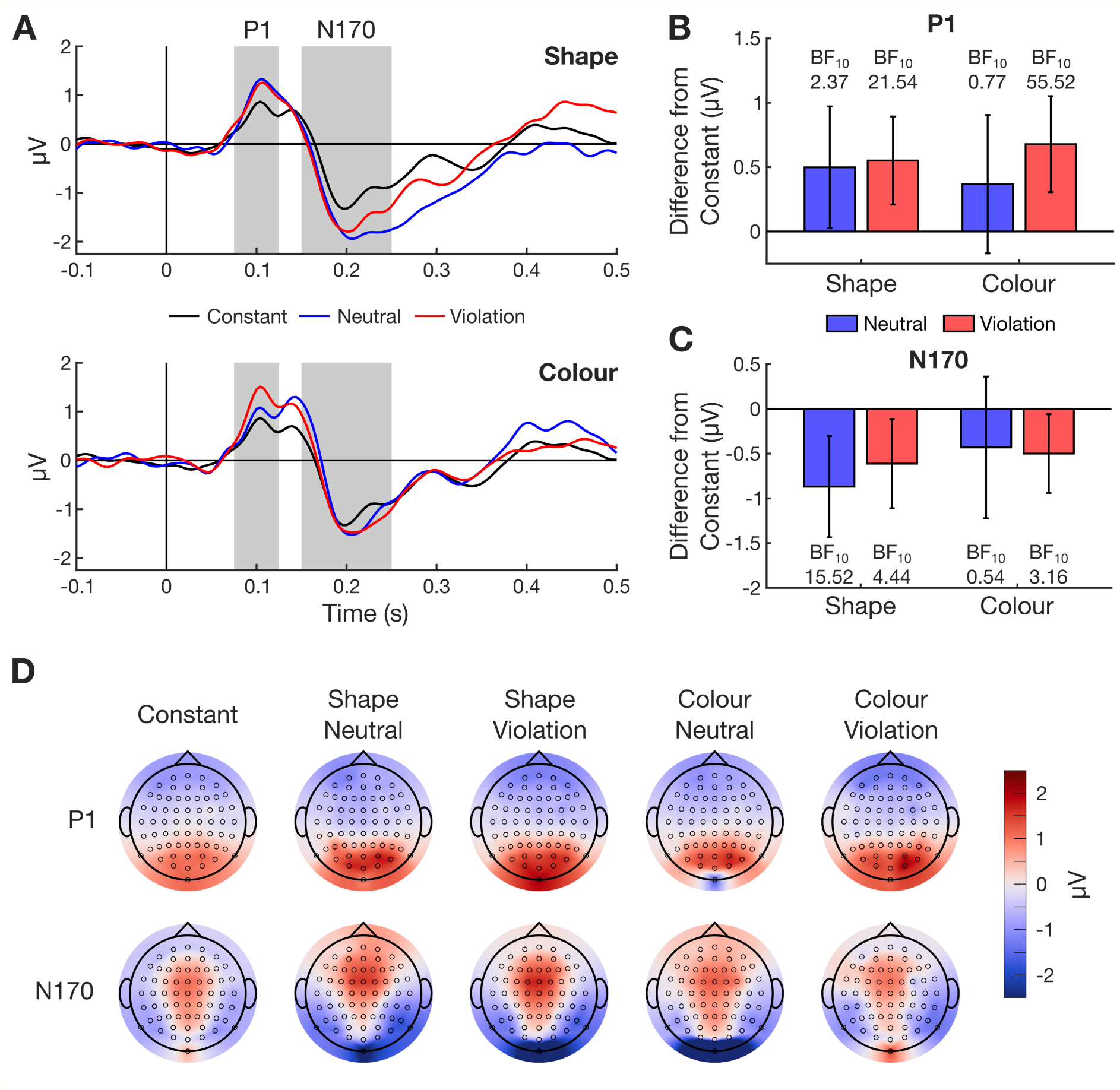
Univariate ERP results. **A)** Grand-average event-related potentials evoked from the fourth image step in each condition. Evoked responses were averaged over P7, P8, P9, P10, PO7, PO8, O1, and O2. The P1 and N170 time windows are filled in grey. **B-C)** Difference in peak amplitude between the constant condition and each change trajectory condition for the P1 and N170 components, respectively. Error bars denote 95% confidence intervals (*N* = 32). Bayes factors quantify the amount of evidence favouring the alternative hypothesis (one-tailed). **D)** Whole-scalp topographic plots within each time window across conditions.

We then tested whether the peak P1 and N170 component amplitudes (shown in Figure 2A-B) differed between change trajectory types and attributes. For the P1, we found evidence for no main effect of both change trajectory type (BF_incl_ = 0.24) and attribute (BF_incl_ = 0.16). Furthermore, these factors did not interact (BF_incl_ = 0.06). The same pattern was observed for the N170 component. Specifically, we found null hypothesis support for a main effect of both change trajectory type (BF_incl_ = 0.2) and attribute (BF_incl_ = 0.27), with these factors showing no interaction (BF_incl_ = 0.08). Thus, while the univariate amplitudes of ERP responses were affected by whether a stimulus attribute changed during the fourth trajectory step, this effect appeared largely agnostic towards how a trajectory was changing.

### Decoding Attribute-Specific Change Context Signalling

Next, we tested whether we could decode the change trajectory that fourth-step stimuli were shown within, despite the above univariate analysis finding no ERP differences between change trajectory types. Crucially, we did this in an attribute-specific manner, using otherwise identical stimuli between the conditions being compared (see Figure 1A). Thus, any decoded signal must have reflected modulatory effects driven by stimulus *context* (neutral vs. violation)—i.e., how surprising an attribute onset was.

Indeed, we were able to decode change context for both shape and colour, with both decoding traces containing qualitatively similar components, particularly from ∼250 ms post stimulus onset (Figure 3A). Interestingly, our searchlight analysis suggested that these multivariate differences were evident throughout the scalp, with classification accuracies being greatest within central electrodes rather than posterior ones—as one might expect under strict interpretations of predictive coding (Friston, 2005). Moreover, despite change context decoding traces being larger, and onsetting earlier, for shape than colour, only seven post-stimulus time points showed at least ‘moderate’ evidence (BF_10_ > 3) for a difference in classification accuracy between the two attributes. In fact, the large majority of time points exhibited evidence for no differences in decodability between the attributes (Figure 3C).

**Figure 3.**
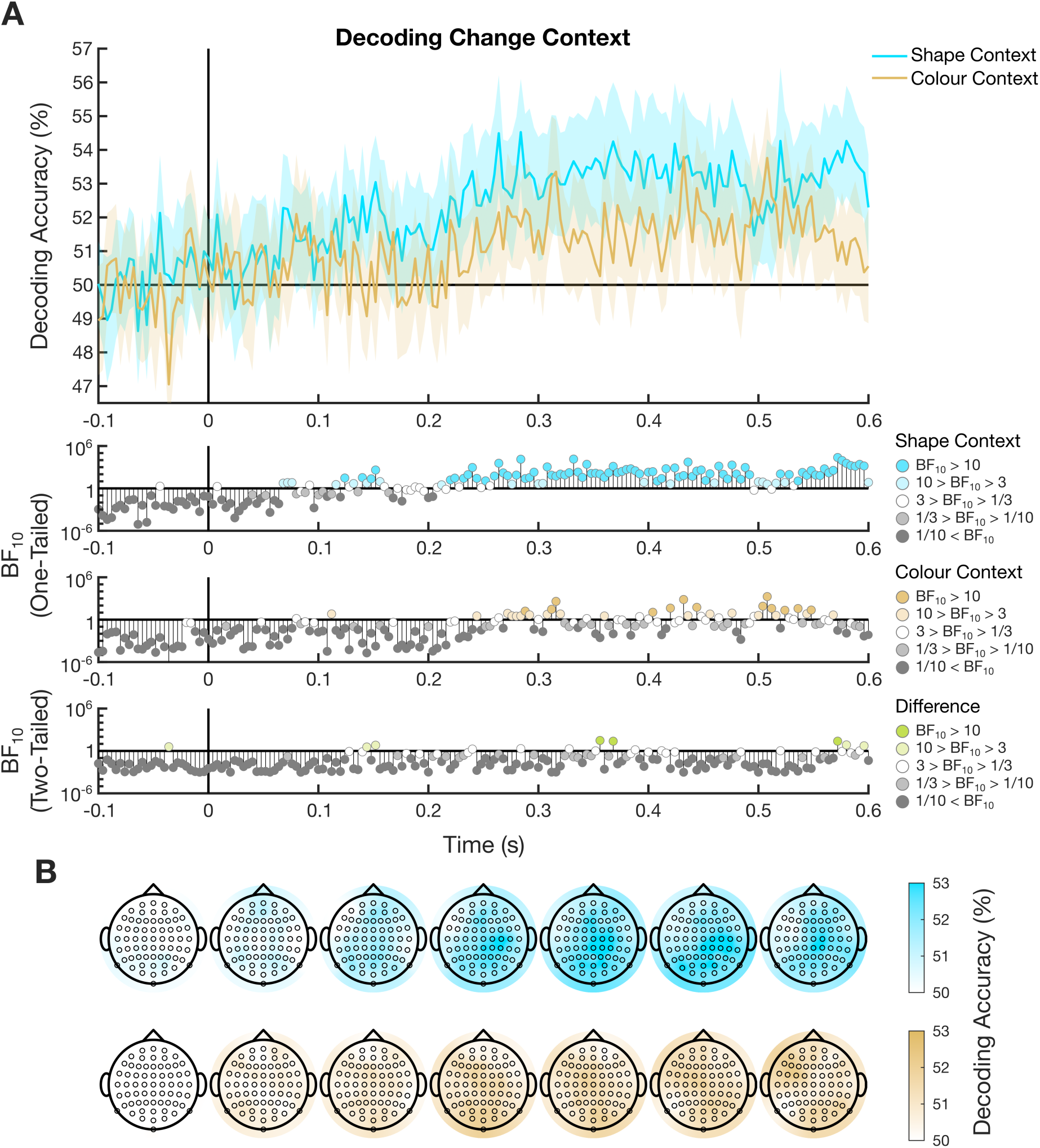
Decoding change context within each attribute. **A)** Change context decoding traces in response to the fourth image step (neutral vs. violation), separated by attributes (shape and colour). Error bars denote 95% confidence intervals (*N* = 32). Stimulus onset is denoted by the solid vertical line, and chance classification accuracy (50%) is denoted by the solid horizontal line. One-tailed Bayes factors quantifying evidence for above-chance classification accuracy across time points are shown within the below subplots. Two-tailed Bayes factors quantifying evidence for a difference in classification accuracy between the two time series are shown in green. **B)** Searchlight results averaged over 100-ms time windows for the shape and colour context comparisons plotted within **A**.

### Cross-Decoding Change Context Between Attributes

After establishing scalp-wide change context signalling evoked from both shape and colour sequences, we tested whether these signals shared overlapping representations between attributes using a temporally generalised cross-decoding approach (Figure 4). This found very clear and sustained effects, particularly from ∼250 ms across both the training and testing time points. Crucially, this overlap predominantly clustered off of the TGM diagonals (*p* < 0.05; cluster-level inference), suggesting the presence of a latency shift between attributes (King & Dehaene, 2014). Most of this overlap occurred when classifiers were trained to decode change context on earlier colour time points relative to shape (and later shape time points relative to colour); however, the opposite pattern of results was also observed across a smaller number of time points within the same signal clusters. These data further support the notion of shared surprise signalling across attributes within bound stimulus objects (Jiang et al., 2016; Lowe et al., 2023).

**Figure 4.**
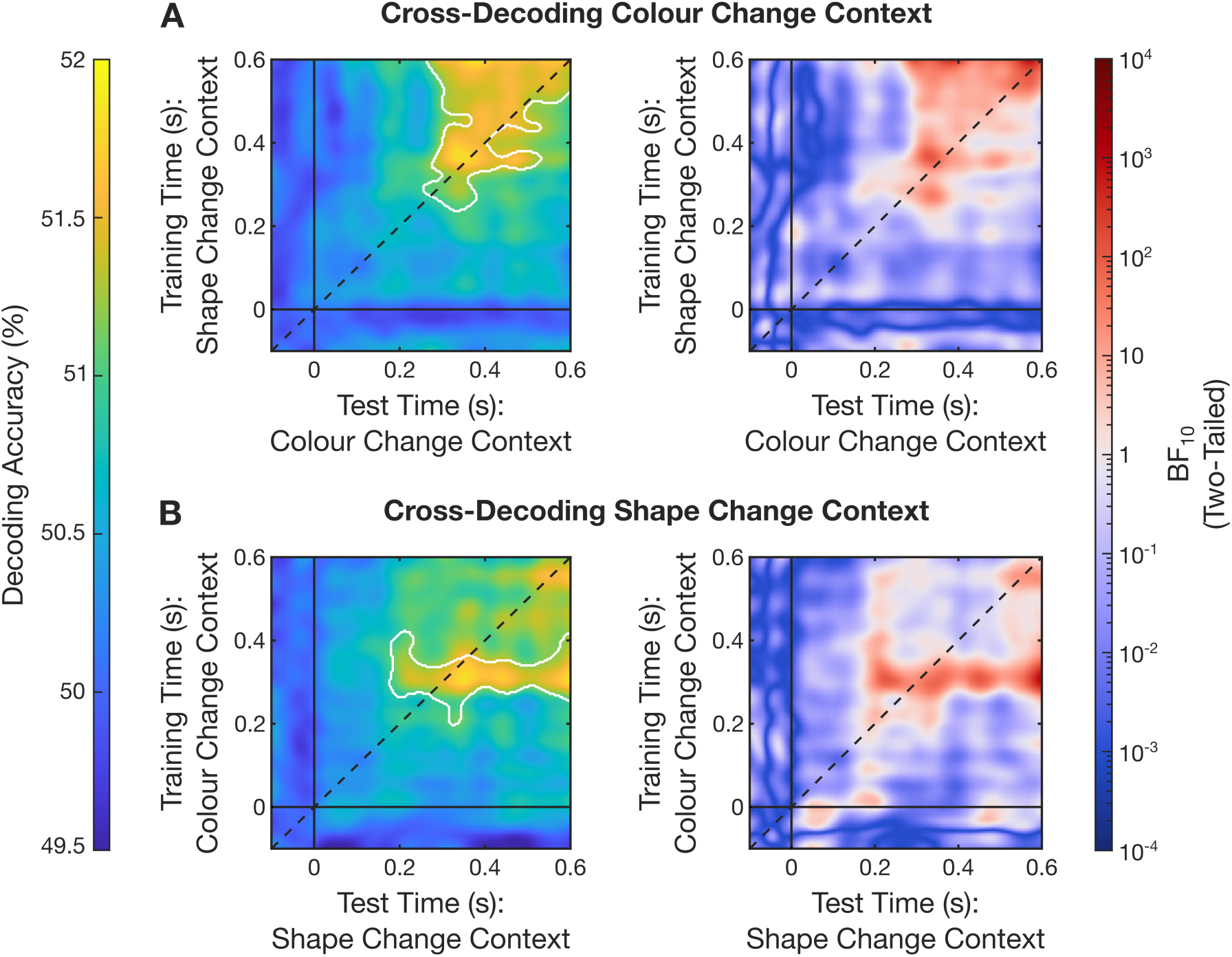
Temporally generalised cross-decoding between attributes. **A)** Classifiers trained to decode change context (neutral vs. violation sequences) from the fourth image step for shape were used to cross-decode change context for colour in a temporally generalised manner. Solid black lines denote stimulus onset in both training and testing time, while the dashed diagonal line marks when training and test times are equivalent. The left matrix depicts cell-wise classification accuracies, with the solid white outlines indicating significantly large clusters of cross-decoded (i.e., shared) signalling with a family-wise error rate of 0.05. The right matrix depicts cell-wise Bayes factors quantifying evidence for two-tailed differences from chance-level (i.e., 50%) classification accuracy. Red cells indicate alternative hypothesis support for a cross-decoded signal, whilst blue cells indicate null hypothesis support. **B)** Classifiers trained to decode change context for colour were used to cross-decode change context for shape. Plotting convention is the same as in **A.** Unsmoothed depictions of the same data can be seen within Figure S2.

## Discussion

The present study investigated whether the timing of visually evoked surprise signals, as recorded by EEG, vary across stimulus attributes. Specifically, we manipulated the context of a shape or colour change, such that it either violated or conformed to an established trajectory (Figure 1). Violation changes were assumed to be surprising, while neutral changes were assumed to be unsurprising. Despite finding predominantly null support for surprise-related ERP component differences (Figure 2), we successfully decoded whether an attribute change was presented within a surprising context from the multivariate patterns of neural activity (Figure 3). Crucially, these change context signals shared overlapping representations between attributes, albeit at different latencies (Figure 4). These findings build upon previous work showing that prediction error responses are shared across attributes (Jiang et al., 2016), and further support the notion of a temporal hierarchy in visual surprise (Lowe et al., 2023).

Since Mishkin and Ungerleider’s (1982) demarcation of the visual system into an object-sensitive ventral stream and spatially-sensitive dorsal stream, there exists a plethora of work highlighting the functionally specialised organisation of visual cortex. Not only are different attributes encoded within different cortical locations (Barrett et al., 2001; Goodale et al., 1991; Haxby et al., 1991; Mishkin et al., 1983; Seymour et al., 2010; Zeki et al., 1991), they are also encoded at different latencies (Arnold, 2005; Clifford et al., 2003, 2004; Holcombe, 2009; Moutoussis & Zeki, 1997b, 1997a). To date, only a handful of studies have investigated how prediction error signalling interacts with the specific attributes that violate an expectation (Jiang et al., 2016; Lowe et al., 2023; Robinson, Woods, et al., 2020; Stefanics et al., 2018, 2019), with these being principally concerned with localising the origins of prediction error in particular brain regions. Our work furthers these insights, emphasising the importance of the temporal domain in prediction error signalling.

The decoded time series for both shape and colour surprise signalling were highly correlated, sharing many of the same peaks and troughs (Figure 3). It could be argued that signalling was more intense and onset earlier for shape compared to colour; however, because this difference was only qualitative, we derive no strong conclusions from this comparison. Interestingly, the qualitative searchlight results implicated responses across the entire scalp in signalling surprise. This is counter to the notion that prediction errors are signalled within the sensory cortex (Friston, 2005) but agrees with more recent empirical work (Westerberg et al., 2025).

Our main assertion of a generalised prediction error response stems from our ability to cross-decode change context (i.e., how surprising a stimulus was) between attributes from ∼250 ms post stimulus onset (Figure 4). Specifically, classifiers trained on colour change context could decode shape change context (and vice versa) across multiple time points post stimulus onset. Crucially, this overlap predominantly clustered off of the TGM diagonals (where training and test times were equivalent), strongly implying latency shifts in an otherwise shared response (see King & Dehaene, 2014), and neatly replicating our previous work with orientation and size expectations (Lowe et al., 2023). Crucially, the shape and colour changes manipulated here cannot have been conflated with a third, higher-order attribute. This strongly suggests that visual surprise may be a domain-general signal that conforms to an attribute-dependent temporal hierarchy.

At first glance, the nature of this latency shift is somewhat confusing due to both cluster boundaries crossing the TGM diagonals. This could suggest a nonlinear temporal structure in response overlap between attributes, whereby the initial components of the generalised surprise signal are evoked earlier for shape than colour, while the later components have the opposite temporal structure. However, we make this statement with caution, given that our cluster-level inference approach lacks the appropriate spatial specificity for this claim (Friston et al., 1996). This said, similar conclusions can be drawn from the cell-wise Bayes factors that provide ‘extreme support’ for the alternative hypothesis (i.e., BF_10_ > 100; Lee & Wagenmakers, 2014) on both sides of the diagonals (even when the TGMs are left unsmoothed; see Figure S2). As such, this interpretation is possibly not without merit; however, speculating on what this non-linearity might mean for a surprise signal is difficult.

An alternative, and more conservative, approach towards interpreting the latency shifts between attributes is to only consider each cluster’s centroid relative to the TGM diagonal. This heuristic leads to a much simpler inference, where the shared visual surprise response was evoked earlier for colour compared to shape. While less nuanced, this interpretation is compelling given that previous psychophysics work has suggested that colour processing precedes that of form (Moutoussis & Zeki, 1997b). As such, it is fair to assume that a shared surprise response between these attributes would conform to the same temporal hierarchy. Regardless of which interpretation is adopted, it is clear from these data that prediction error signalling shares overlapping responses between attributes when accounting for differences in signal latencies.

This notion aligns well with previous fMRI work by Jiang et al. (2016). There, the authors proposed that prediction error signalling within one functionally segregated region of cortex will ‘spread’ throughout the visual system when viewing a bound object comprised of multiple attributes—even if the other attributes conformed to expectations. Said differently, the entirety of an object is rendered surprising if just one of its attributes violates an expectation. Indeed, given that we used bound stimuli within the present study (Figure 1), it is conceivable that the overlap in neural representation shown in Figure 4 reflects this spreading of prediction error throughout the visual system. In other words, a violation concerning stimulus colour (or shape) caused prediction error signalling for *both* colour and shape. This would explain why classifiers trained to decode change context from one attribute could decode the same information from the other. Here, we have extended Jiang et al.’s model by highlighting that the timing of this overlap likely conforms to a temporal hierarchy.

The notion of generic prediction error responses is difficult to reconcile with classical predictive coding as described by Friston (2005, 2010). In short, this model asserts that all evoked responses signal the mismatch between a sensory prediction (expectation) and input. Because of this, surprise responses are assumed to be attribute-specific and onset during the initial feedforward visual sweep (i.e., earlier than ∼150 ms post-stimulus onset). We failed to observe this pattern of results outside of eight (somewhat sparsely located) time points within the shape change context comparison (Figure 3A).

Instead, these data align more closely with Press et al.’s (2020) opposing process theory of perceptual inference and learning. Here, the feedforward sweep is initially biased towards representing expected input, with prediction error being signalled after ∼150 ms to increase the salience of unexpected inputs and drive learning. This would explain the relatively late onset of our decoded surprise signals (which were undeniable from ∼250 ms post stimulus onset). In extending Press et al.’s theory, our data suggest that this later prediction error signalling process may either operate at the bound object level, or is at least shared across attributes.

Turning now to our univariate results, despite finding clear ERP component differences between the constant sequence condition and almost all change sequence conditions (Figure 2A-C), our Bayesian ANOVAs found null hypothesis support for all main effects and interactions when comparing P1 and N170 amplitudes between change context types (neutral vs. violation) and attributes (shape vs. colour). These results are worth highlighting because they speak to a growing critique within the predictive coding literature concerning how classical surprise responses are not observed after controlling for cortical adaptation (Feuerriegel, 2024; Feuerriegel et al., 2021; Solomon et al., 2021; Westerberg et al., 2025).

To elaborate, a core tenet of predictive coding is that prediction error signalling is excitatory whilst expectation signalling is inhibitory (Bastos et al., 2012; Friston, 2005, 2010; Walsh et al., 2020). Surprising stimulus input should therefore increase the amplitude of evoked responses relative to unsurprising input. However, these effects are difficult to delineate from adaptation if a stimulus is deemed ‘unsurprising’ purely because it is a repeat of another shown moments earlier. This is because when the visual system is repeatedly stimulated by the same stimulus attribute (e.g., the same colour or shape), functionally specialised neurons coding that attribute will exhibit a dampened response profile due to neural fatigue (Feuerriegel, 2024). Therefore, any comparisons between evoked responses involving the constant sequence condition cannot be used as evidence for prediction error signalling, as they cannot account for this confound. This was partially why constant sequence trials were omitted from our decoding analyses.

By contrast, the comparisons between neutral and violation change sequences inherently controlled for adaptation-related confounds because both conditions evoked neural responses to attribute *changes* (rather than repeats; Figure 1A). As mentioned, Bayes factors here supported the null hypothesis, suggesting that there was no effect of surprise (or attribute), which is an interesting deviation from other studies with highly comparable paradigms (Baker et al., 2021, 2022; Johnston et al., 2017; Robinson, Breakspear, et al., 2020; Robinson, Woods, et al., 2020). A reason for this departure may have been our use of a neutral control condition, as opposed to a condition wherein expectations were confirmed, given that surprise and expectation effects may reflect different mechanisms (see Feuerriegel et al., 2021). Alternatively, it may very well be that sensory responses of visual cortical neurons are not prediction errors (Westerberg et al., 2025).

## Conclusion

In sum, we found overlapping surprise responses evoked by both the shape and colour of a bound stimulus object, albeit at distinct latencies. This suggests that visual surprise is (at least partially) a domain-general response that conforms to an attribute-specific temporal hierarchy. Future work delineating the nature of prediction error signalling should therefore account for the specific attributes violating an expectation, particularly concerning the latency of evoked responses.

## Acknowledgements

We would like to acknowledge Turrbal and Jagera peoples, who are the traditional owners of the land on which these data were collected. This work was supported by an Australian Government Research Training Program Scholarship awarded to BGL whilst he was a student at QUT.

## Supplementary Materials

### Temporal Shift in Event Triggers During Preprocessing

As mentioned within the main text, we temporally shifted the timing of the epoched data. This was because trigger codes, signalling the onset of events, were being sent two frames before (33 ms, 60 Hz display) stimulus onset. This is evident within Figure S1, which shows that when the timing of our epoched data were shifted by -33 ms, canonical event-related potential (ERP) components occur within their typical time window.

**Figure S1.**
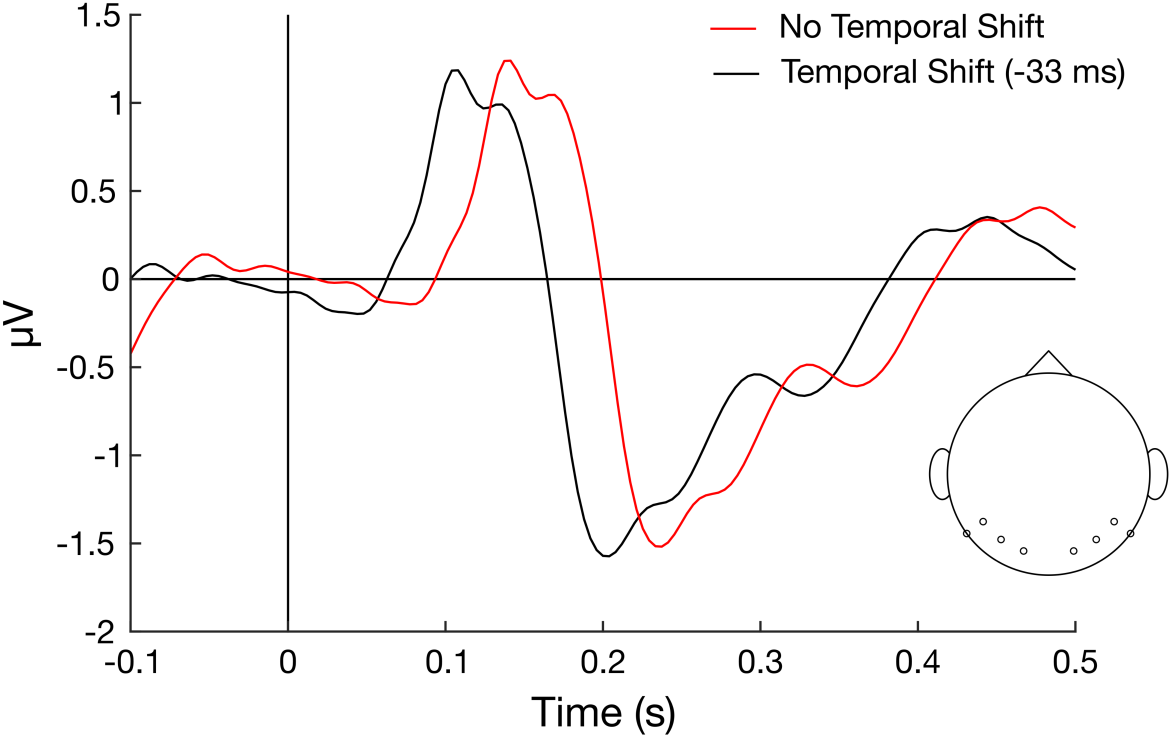
Temporal shift rationale. Posterior event-related potentials evoked from the fourth image step (collapsed across conditions) with and without a -33 ms temporal shift to account for early event trigger times during recording. Notice that without the temporal shift, the P1 and N170 peaks appear substantially later than usual. The circles on the head picture indicate electrodes from which data for these waveforms were collected (O1, O2, PO7, PO8, P7, P8, P9, and P10).

### Unsmoothed Cross-Decoding Results

To ensure that our cross-decoding results were not driven by data smoothing, we repeated the same analysis on unsmoothed data, which yielded results that mirrored those reported within the main text (Figure S2). Specifically, cell-wise Bayes factors for the alternative hypothesis found extreme evidence for change context cross-decoding between attributes across both sides of the TGM diagonals (BF_10_ > 100). We also performed cluster-level inference on these data (*p* < 0.05), which showed similar results. Namely, there were significantly large decoding accuracy clusters level the diagonal when cross-decoding shape change context from colour. By contrast, significant decoding accuracy clusters were predominantly located above the diagonal when cross-decoding colour change context from shape—though two clusters were found below the diagonal. Note that performing cluster-level inference on unsmoothed data is not standard practice, and these results have just been presented for the reader’s interest.

**Figure S2.**
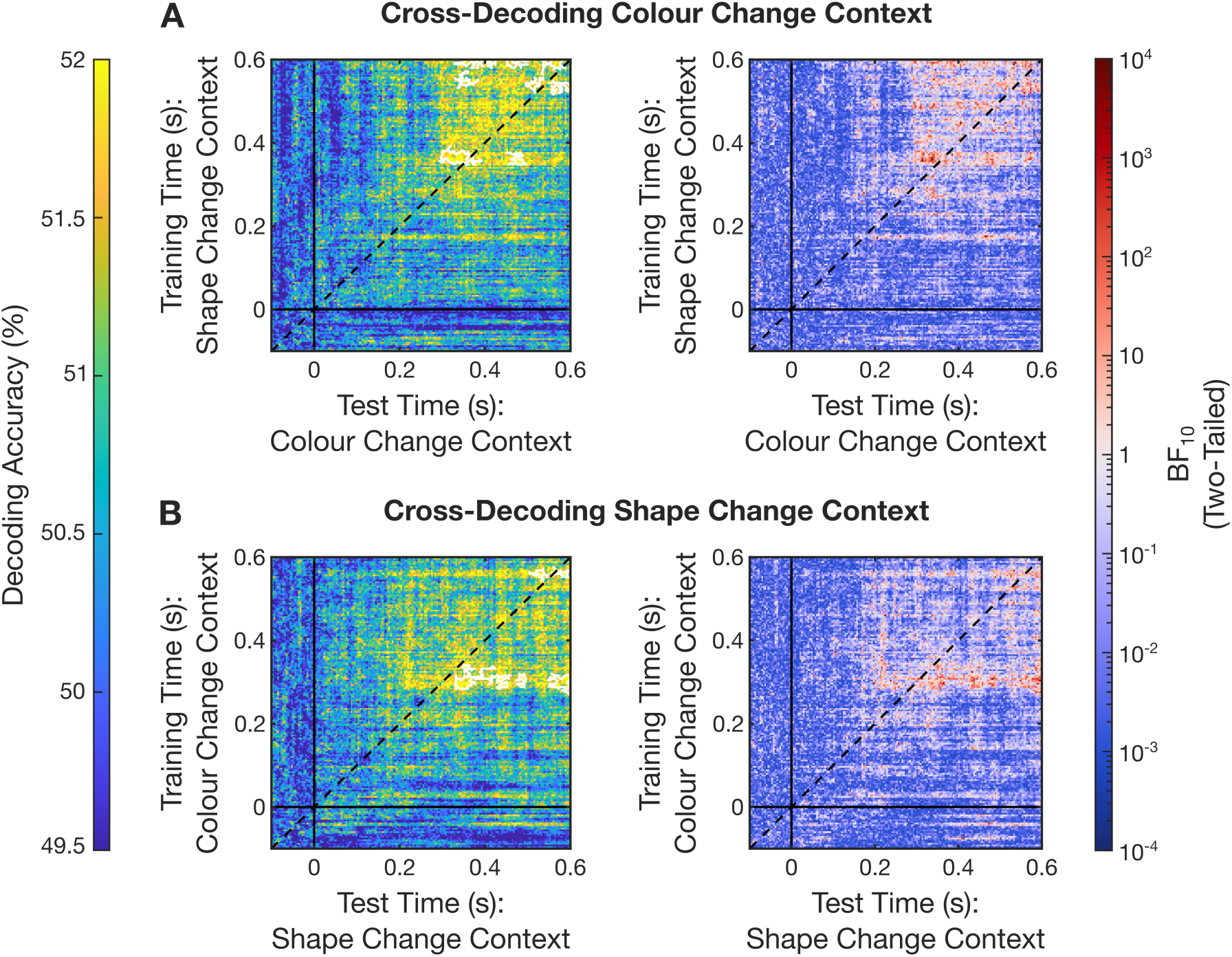
Temporally generalised cross-decoding between attributes (unsmoothed). **A)** Classifiers trained to decode change context (neutral vs. violation sequences) from the fourth image step for shape were used to cross-decode change context for colour in a temporally generalised manner. Solid black lines denote stimulus onset in both training and testing time, while the dashed diagonal line marks when training and test times are equivalent. The left matrix depicts cell-wise classification accuracies, with the solid white outlines indicating significantly large clusters of cross-decoded (i.e., shared) signalling with a family-wise error rate of 0.05. The right matrix depicts cell-wise Bayes factors quantifying evidence for two-tailed difference from chance-level (i.e., 50%) classification accuracy. Red cells indicate alternative hypothesis support for a cross-decoded signal, whilst blue cells indicate null hypothesis support. **B)** Classifiers trained to decode change context for colour were used to cross-decode change context for shape. Plotting convention is the same as in **A.** Smoothed depictions of the same data, which are more appropriate for cluster-level analysis, can be seen within Figure 4 of the main text.

### Orientation Surprise Decoding

As mentioned within the main text, we intended to investigate the dynamics of orientation surprise signalling in addition to that of shape and colour. This, however, was omitted from the main analyses because the attribute lacked a neutral change condition. Instead, we had planned to compare responses between violation trials and matched control trials (Figure S3A). Note that orientation changes were designed so that clockwise rotations increased in 20° increments of polar angle, whilst counter-clockwise rotations decreased in 15° increments (counter-balanced across participants). Thus, the final image step’s orientation (step 4) was never a repeat of that shown two steps earlier (step 2), even when the sequence was violated, controlling for repetition suppression.

Like with shape and colour, we decoded change context for orientation, this time using constant sequences instead of attribute-specific neutral ones. Again, this was done using 96-fold cross-validation with a subset of constant trials that matched the stimuli used for orientation violations whilst having minimal differences in onset times between conditions (Figure S3B). The decoding trace was virtually identical to that found in our previous work using the same comparison (Lowe et al., 2023). Qualitative searchlight analysis emphasised the importance of central and posterior regions.

**Figure S3.**
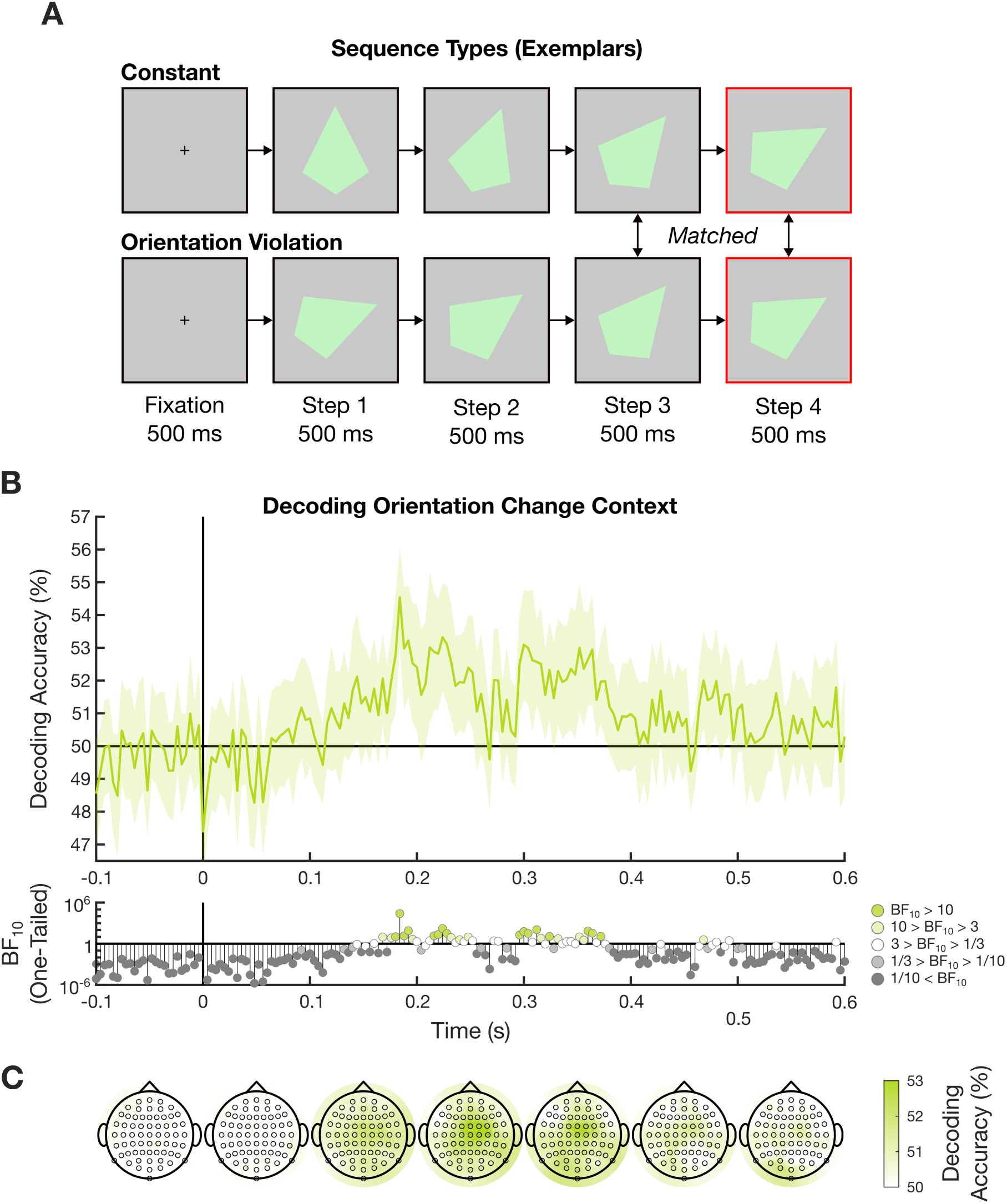
Decoding orientation change context. **A)** Exemplars of each sequence type. Note that the final two steps are identical across the violation and constant conditions. Any differences in evoked responses from Step 4 (red outline) must have therefore reflected the preceding change context. **B)** Change context decoding traces in response to the fourth image step (constant vs. violation). Error bars denote 95% confidence intervals (*N* = 32). Stimulus onset is denoted by the solid vertical line and chance classification accuracy (50%) is denoted by the solid horizontal line. One-tailed Bayes factors quantifying evidence for an above-chance difference across time points are shown within the below subplots. **C)** Searchlight results averaged over 100-ms time windows for the same comparison plotted within **B**.

### Orientation Surprise Cross-Decoding

Having established a surprise response for orientation, we then used the classifiers trained to decode change context for this attribute to cross-decode the same signal for both colour and shape (Figure S4). Admittedly, it is not straightforward to interpret the outcomes of this approach because of the different control conditions used between the training (i.e., constant sequences) and test attributes (i.e., neutral sequences). Nevertheless, we still found clear overlapping responses between orientation and shape. Interestingly, this occurred even earlier than the overlap between shape and colour reported within the main text. Cell-wise Bayes factors suggested that there might have been later overlap between orientation and colour, but this cluster was too small to be deemed significant using cluster-level inference.

We are careful not to draw strong conclusions about domain-general surprise signalling from these data because of the differences in control condition sequence types between attributes.

**Figure S4.**
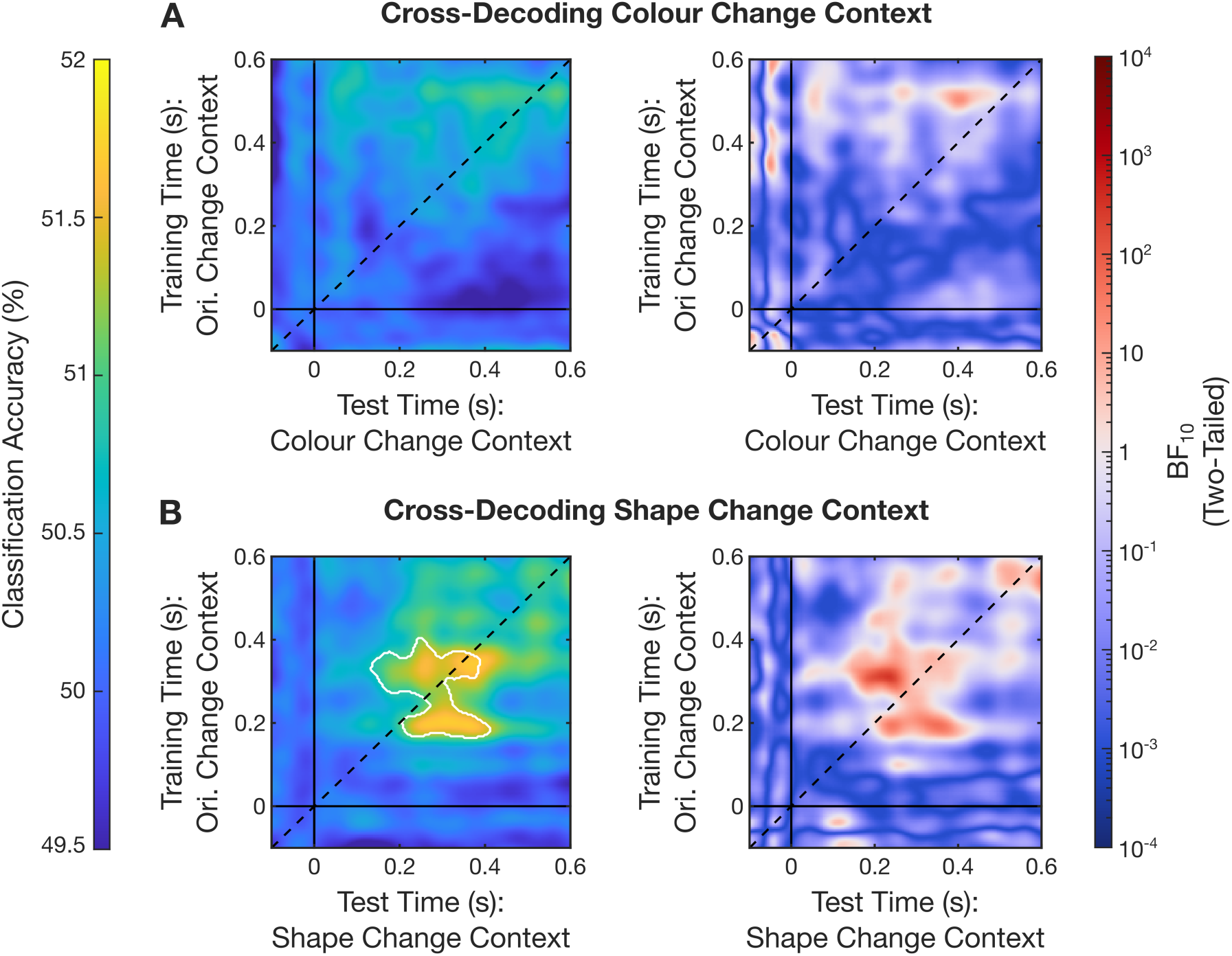
Temporally generalised cross-decoding between attributes from orientation change sequence data. **A)** Classifiers trained to decode orientation change context (constant vs. violation sequences) were used to cross-decode change context for colour in a temporally generalised manner. Solid black lines denote stimulus onset in both training and test time, while the dashed diagonal line marks when training and test times are equivalent. The left matrix depicts cell-wise classification accuracies, with the solid white outlines indicating significantly large clusters of cross-decoded (i.e., shared) signalling with a family-wise error rate of 0.05 (none were found). The right matrix depicts cell-wise Bayes factors quantifying evidence for two-tailed differences from chance-level (i.e., 50%) classification accuracy. Red cells indicate alternative hypothesis support for a cross-decoded signal, whilst blue cells indicate null hypothesis support **B)** The same method applied to shape. Plotting convention is the same as in **A.**

## References

Amado, C., & Kovács, G. (2016). Does surprise enhancement or repetition suppression explain visual mismatch negativity? European Journal of Neuroscience, 43(12), 1590–1600. 10.1111/ejn.13263

Arnold, D. H. (2005). Perceptual pairing of colour and motion. Vision Research, 45(24), 3015–3026. 10.1016/j.visres.2005.06.031

Baker, K. S., Pegna, A. J., Yamamoto, N., & Johnston, P. (2021). Attention and prediction modulations in expected and unexpected visuospatial trajectories. PLOS ONE, 16(10), e0242753. 10.1371/journal.pone.0242753

Baker, K. S., Yamamoto, N., Pegna, A. J., & Johnston, P. (2022). Violated expectations for spatial and feature attributes of visual trajectories modulate event-related potential amplitudes across the visual processing hierarchy. Biological Psychology, 174, 108422. 10.1016/j.biopsycho.2022.108422

Barrett, N. A., Large, M. M., Smith, G. L., Michie, P. T., Karayanidis, F., Kavanagh, D. J., Fawdry, R., Henderson, D., & O’Sullivan, B. T. (2001). Human cortical processing of colour and pattern. Human Brain Mapping, 13(4), 213–225. 10.1002/hbm.1034

Bastos, A. M., Usrey, W. M., Adams, R. A., Mangun, G. R., Fries, P., & Friston, K. J. (2012). Canonical Microcircuits for Predictive Coding. Neuron, 76(4), 695–711. 10.1016/j.neuron.2012.10.038

Clifford, C. W. G., Arnold, D. H., & Pearson, J. (2003). A paradox of temporal perception revealed by a stimulus oscillating in colour and orientation. Vision Research, 43(21), 2245–2253. 10.1016/S0042-6989(03)00120-2

Clifford, C. W. G., Holcombe, A. O., & Pearson, J. (2004). Rapid global form binding with loss of associated colors. Journal of Vision, 4(12), 1090–1101. 10.1167/4.12.8

Feuerriegel, D. (2024). Adaptation in the visual system: Networked fatigue or suppressed prediction error signalling? Cortex, 177, 302–320. 10.1016/j.cortex.2024.06.003

Feuerriegel, D., Vogels, R., & Kovács, G. (2021). Evaluating the evidence for expectation suppression in the visual system. Neuroscience & Biobehavioral Reviews, 126, 368–381. 10.1016/j.neubiorev.2021.04.002

Friston, K. (2005). A theory of cortical responses. Philosophical Transactions of the Royal Society B: Biological Sciences, 360(1456), 815–836. 10.1098/rstb.2005.1622

Friston, K. (2010). The free-energy principle: A unified brain theory? Nature Reviews Neuroscience, 11(2), 127–138. 10.1038/nrn2787

Friston, K. J., Holmes, A., Poline, J.-B., Price, C. J., & Frith, C. D. (1996). Detecting Activations in PET and fMRI: Levels of Inference and Power. NeuroImage, 4(3), 223–235. 10.1006/nimg.1996.0074

Goodale, M. A., Milner, A. D., Jakobson, L. S., & Carey, D. P. (1991). A neurological dissociation between perceiving objects and grasping them. Nature, 349(6305), 154–156. 10.1038/349154a0

Gramfort, A., Luessi, M., Larson, E., Engemann, D. A., Strohmeier, D., Brodbeck, C., Goj, R., Jas, M., Brooks, T., Parkkonen, L., & Hämäläinen, M. (2013). MEG and EEG data analysis with MNE-Python. Frontiers in Neuroscience, 7. 10.3389/fnins.2013.00267

Grotheer, M., & Kovács, G. (2016). Can predictive coding explain repetition suppression? Cortex, 80, 113–124. 10.1016/j.cortex.2015.11.027

Haxby, J. V., Grady, C. L., Horwitz, B., Ungerleider, L. G., Mishkin, M., Carson, R. E., Herscovitch, P., Schapiro, M. B., & Rapoport, S. I. (1991). Dissociation of object and spatial visual processing pathways in human extrastriate cortex. Proceedings of the National Academy of Sciences, 88(5), 1621–1625. 10.1073/pnas.88.5.1621

Holcombe, A. O. (2009). Seeing slow and seeing fast: Two limits on perception. Trends in Cognitive Sciences, 13(5), 216–221. 10.1016/j.tics.2009.02.005

Jiang, J., Summerfield, C., & Egner, T. (2016). Visual Prediction Error Spreads Across Object Features in Human Visual Cortex. Journal of Neuroscience, 36(50), 12746–12763. 10.1523/JNEUROSCI.1546-16.2016

Johnston, P., Robinson, J., Kokkinakis, A., Ridgeway, S., Simpson, M., Johnson, S., Kaufman, J., & Young, A. W. (2017). Temporal and spatial localization of prediction-error signals in the visual brain. Biological Psychology, 125, 45–57. 10.1016/j.biopsycho.2017.02.004

King, J.-R., & Dehaene, S. (2014). Characterizing the dynamics of mental representations: The temporal generalization method. Trends in Cognitive Sciences, 18(4), 203–210. 10.1016/j.tics.2014.01.002

Kriegeskorte, N., Goebel, R., & Bandettini, P. (2006). Information-based functional brain mapping. Proceedings of the National Academy of Sciences of the United States of America, 103(10), 3863–3868. 10.1073/pnas.0600244103

Lee, M. D., & Wagenmakers, E.-J. (2014). Bayesian Cognitive Modeling: A Practical Course. Cambridge University Press.

Lowe, B. G., Robinson, J. E., Yamamoto, N., Hogendoorn, H., & Johnston, P. (2023). Same but different: The latency of a shared expectation signal interacts with stimulus attributes. Cortex, 168, 143–156. 10.1016/j.cortex.2023.08.004

Maris, E., & Oostenveld, R. (2007). Nonparametric statistical testing of EEG- and MEG-data. Journal of Neuroscience Methods, 164(1), 177–190. 10.1016/j.jneumeth.2007.03.024

Mishkin, M., & Ungerleider, L. G. (1982). Contribution of striate inputs to the visuospatial functions of parieto-preoccipital cortex in monkeys. Behavioural Brain Research, 6(1), 57–77. 10.1016/0166-4328(82)90081-X

Mishkin, M., Ungerleider, L. G., & Macko, K. A. (1983). Object vision and spatial vision: Two cortical pathways. Trends in Neurosciences, 6, 414–417. 10.1016/0166-2236(83)90190-X

Morey, R., & Rouder, J. (2023). BayesFactor: Computation of Bayes Factors for Common Designs (Version 0.9.12-4.6) [Computer software]. https://github.com/richarddmorey/bayesfactor

Moutoussis, K., & Zeki, S. (1997a). A direct demonstration of perceptual asynchrony in vision. Proceedings of the Royal Society of London. Series B: Biological Sciences, 264(1380), 393–399. 10.1098/rspb.1997.0056

Moutoussis, K., & Zeki, S. (1997b). Functional segregation and temporal hierarchy of the visual perceptive systems. Proceedings of the Royal Society of London. Series B: Biological Sciences. 10.1098/rspb.1997.0196

Peirce, J., Gray, J. R., Simpson, S., MacAskill, M., Höchenberger, R., Sogo, H., Kastman, E., & Lindeløv, J. K. (2019). PsychoPy2: Experiments in behavior made easy. Behavior Research Methods, 51(1), 195–203. 10.3758/s13428-018-01193-y

Press, C., Kok, P., & Yon, D. (2020). The Perceptual Prediction Paradox. Trends in Cognitive Sciences, 24(1), 13–24. 10.1016/j.tics.2019.11.003

Robinson, J. E., Breakspear, M., Young, A. W., & Johnston, P. J. (2020). Dose- dependent modulation of the visually evoked N1/N170 by perceptual surprise: A clear demonstration of prediction-error signalling. European Journal of Neuroscience, 52(11), 4442–4452. 10.1111/ejn.13920

Robinson, J. E., Woods, W., Leung, S., Kaufman, J., Breakspear, M., Young, A. W., & Johnston, P. J. (2020). Prediction-error signals to violated expectations about person identity and head orientation are doubly-dissociated across dorsal and ventral visual stream regions. NeuroImage, 206, 116325. 10.1016/j.neuroimage.2019.116325

Rouder, J. N., Morey, R. D., Speckman, P. L., & Province, J. M. (2012). Default Bayes factors for ANOVA designs. Journal of Mathematical Psychology, 56(5), 356–374. 10.1016/j.jmp.2012.08.001

Rouder, J. N., Speckman, P. L., Sun, D., Morey, R. D., & Iverson, G. (2009). Bayesian t tests for accepting and rejecting the null hypothesis. Psychonomic Bulletin & Review, 16(2), 225–237. 10.3758/PBR.16.2.225

Schneider, G. E. (1969). Two Visual Systems. Science, 163(3870), 895–902.

Seymour, K., Clifford, C. W. G., Logothetis, N. K., & Bartels, A. (2010). Coding and Binding of Color and Form in Visual Cortex. Cerebral Cortex, 20(8), 1946– 1954. 10.1093/cercor/bhp265

Solomon, S. S., Tang, H., Sussman, E., & Kohn, A. (2021). Limited Evidence for Sensory Prediction Error Responses in Visual Cortex of Macaques and Humans. Cerebral Cortex, 31(6), 3136–3152. 10.1093/cercor/bhab014

Stefanics, G., Heinzle, J., Horváth, A. A., & Stephan, K. E. (2018). Visual Mismatch and Predictive Coding: A Computational Single-Trial ERP Study. Journal of Neuroscience, 38(16), 4020–4030. 10.1523/JNEUROSCI.3365-17.2018

Stefanics, G., Stephan, K. E., & Heinzle, J. (2019). Feature-specific prediction errors for visual mismatch. NeuroImage, 196, 142–151. 10.1016/j.neuroimage.2019.04.020

Teichmann, L., Moerel, D., Baker, C., & Grootswagers, T. (2022). An Empirically Driven Guide on Using Bayes Factors for M/EEG Decoding. Aperture Neuro, 2, 1–10. 10.52294/ApertureNeuro.2022.2.MAOC6465

Walsh, K. S., McGovern, D. P., Clark, A., & O’Connell, R. G. (2020). Evaluating the neurophysiological evidence for predictive processing as a model of perception. Annals of the New York Academy of Sciences, 1464(1), 242–268. 10.1111/nyas.14321

Westerberg, J. A., Xiong, Y. S., Sennesh, E., Nejat, H., Ricci, D., Durand, S., Hardcastle, B., Cabasco, H., Belski, H., Bawany, A., Gillis, R., Loeffler, H., Peene, C. R., Han, W., Nguyen, K., Ha, V., Johnson, T., Grasso, C., Young, A., … Bastos, A. M. (2025). Sensory responses of visual cortical neurons are not prediction errors (p. 2024.10.02.616378). bioRxiv. 10.1101/2024.10.02.616378

Zeki, S., Watson, J. D., Lueck, C. J., Friston, K. J., Kennard, C., & Frackowiak, R. S. (1991). A direct demonstration of functional specialization in human visual cortex. Journal of Neuroscience, 11(3), 641–649. 10.1523/JNEUROSCI.11-03-00641.1991

